# Real-time spectral library matching for sample multiplexed quantitative proteomics

**DOI:** 10.1101/2023.02.08.527705

**Authors:** Chris D. McGann, Will Barshop, Jesse Canterbury, Chuwei Lin, Wassim Gabriel, Mathias Wilhelm, Graeme McAlister, Devin K. Schweppe

## Abstract

Sample multiplexed quantitative proteomics has proved to be a highly versatile means to assay molecular phenotypes. Yet, stochastic precursor selection and precursor co-isolation can dramatically reduce the efficiency of data acquisition and quantitative accuracy. To address this, intelligent data acquisition (IDA) strategies have recently been developed to improve instrument efficiency and quantitative accuracy for both discovery and targeted methods. Towards this end, we sought to develop and implement a new real-time library searching (RTLS) workflow that could enable intelligent scan triggering and peak selection within milliseconds of scan acquisition. To ensure ease of use and general applicability, we built an application to read in diverse spectral libraries and file types from both empirical and predicted spectral libraries. We demonstrate that RTLS methods enable improved quantitation of multiplexed samples, particularly with consideration for quantitation from chimeric fragment spectra. We used RTLS to profile proteome responses to small molecule perturbations and were able to quantify up to 15% more significantly regulated proteins in half the gradient time as traditional methods. Taken together, the development of RTLS expands the IDA toolbox to improve instrument efficiency and quantitative accuracy in sample multiplexed analyses.

## Introduction

Nearly every data-dependent analysis suffers from stochastic precursor selection effects. These effects reduce run-to-run coverage of the proteome, increasing the number of missing values, and limiting quantitative access across the proteome^1^. This problem is particularly challenging for methods that require longer scan or fill times to reach necessary resolutions or sensitivity. For example, the most accurate methods for sample multiplexed quantitative proteomics require tertiary quantitative scans - synchronous precursor selection SPS-MS3 scans^2,3^. Acquisition of these scans leads to multifold increases in the time necessary to quantify proteomics samples at sufficient depth (~8000 proteins).

To combat these challenges, intelligent data acquisition (IDA) methods have been employed to improve both the throughput and quantitative accuracy of sample multiplexed proteomics workflows^4,5^. Initially this work focused on optimizing fragmentation of precursors in real-time with instrument acquisition. Since then, these efforts have expanded to improve targeted^6,7^ and discovery-based proteomic workflows^8^. IDA strategies rely on interpretation and utilization of raw spectral data within a few milliseconds of scan acquisition. This data can then be used to trigger additional scans, optimize scan parameters, or filter low utility spectra^9^.

In previous work, we showed that real-time database searching (RTS) could double instrument acquisition efficiency for whole proteome profiling^9^. The Comet-based^10^ peptide spectral matching enabled real-time scoring to inform the instrument when to collect an SPS-MS3 scan and to target b- and y-ions for SPS selection. While RTS is a powerful means to improve instrument acquisition efficiency, we know from *post hoc* analysis of proteomics datasets that additional scoring metrics can increase the sensitivity of peptide detection^11–14^.

One means to increase peptide spectral matching efficiency is through the use of spectral library peptide matching^15^. Early work with spectral library searching for proteomics relied on the construction of empirically derived spectra to generate libraries using well established workflows such as SpectraST to confidently match peptides based on common score metrics (dot product, cosine score, spectral similarity)^16,17^. Recent advances in deep learning have now contributed multiple pipelines for the *in silico* prediction of peptide spectra^11,18^. Algorithms such as Prosit enable users to predict peptide spectra for whole proteomes (2.6 million peptide spectra for the human proteome) and have recently been extended to incorporate isobaric labelled samples^19,20^. These predicted spectral libraries can then be used to efficiently score new empirical spectra or combined with database searching algorithms to re-score spectra for improved sensitivity^21–23^. Spectral library searching has been shown to be a sensitive and accurate way to identify peptides, especially those of complex spectra such as data-independent acquisition experiments^24–28^. Lessons learned from using spectral libraries in those data-independent acquisition experiments has recently been leveraged to improve spectral library searches for data-dependent acquisition methods. These latest spectral library search algorithms show that while using even a predicted library, there is a sensitivity gain compared to cutting-edge database search methods^29^.

Here, we sought to develop **R**eal-**T**ime spectral **L**ibrary **S**earch (RTLS) for whole proteome, sample multiplexed quantitative proteomics. Previous work established that RTLS could be used for small molecule spectral matching with spectral libraries of a single small molecule, and to improve analysis of cross-linked peptides with a two-spectrum library of diagnostic ions^30^. To enable RTLS for whole proteome work, we needed to expand on these efforts to enable RTLS to process full proteome libraries of up to 2 million spectra in real time with spectral acquisition. To accomplish this, we (1) developed a flexible library processing workflow to enable the use of both predicted and experimental libraries from common proteomics resources, (2) optimized our online scoring for whole proteome sample multiplexed analyses, (3) determined optimal parameters for improved instrument efficiency and quantitative accuracy, and (4) compared the RTLS workflow to traditional acquisition methods using established standards and complex proteomic samples. Using our optimized workflow and methods, RTLS increased instrument acquisition efficiency 2-fold, in keeping with previous IDA methods. In addition, RTLS improved quantitative accuracy for chimeric spectra from whole proteome single shot samples and enabled fast and efficient identification of post-translationally modified peptides. Thus, RTLS has proved to be a useful addition to the IDA toolkit with great potential for sample multiplexed quantitative proteomics and future development.

## Experimental Procedures

### Sample collection and preparation

Human cell lines were grown to confluence in DMEM containing 10% fetal bovine serum and 1% streptomycin/puromycin. Cells were harvested by manual scraping and washed twice with PBS. Cells were syringe lysed in lysis buffer (8M urea, 50mM EPPS pH 8.5, 150mM NaCl, and Roche protease inhibitor tablet) and the resulting lysates were cleared via centrifugation.

*Saccharomyces cerevisiae* (BY4742) was grown YPD cultures to an OD600 of 0.8 then washed twice with PBS, pelleted, and stored at −80 °C until use. Cells were resuspended in lysis buffer (8 M urea, 50 mM EPPS pH 8.5, 150 mM NaCl, Roche protease inhibitor tablet) and lysed by bead beating. After lysis and bead removal, the lysate was centrifuged to remove cellular debris and the supernatant was collected for use.

A two proteome (human and yeast) HyPro standard labeled with TMTpro was prepared as previously described^31^. In brief, HCT116 cells were prepared according to the SL-TMT protocol^32^ and labeled at 1:1 across all channels. *S. cerevisiae* (BY4716) was prepared similarly but with ratios of 0:1:1:1:2:2:2:4:4:4:8:8:8:10:10:10 across the 16 TMTpro reporter ion channels. The HCT116 and *S. cerevisiae* were combined so the final sample was 90% human peptides and 10% yeast peptides (w/w). In the small molecule perturbation studies, A549, H292, or PSC1 cells were treated with 10 μM of a given HDAC inhibitor for 24 hours (belinostat, abexinostat, CUDC-101, vorinostat).

### Mass spectrometry data acquisition methods and analysis

Samples were resuspended in 5% acetonitrile/2% formic acid prior to being loaded onto an in-house pulled C18 (Thermo Accucore, 2.6 Å, 150 μm) 30 cm column. Peptides were eluted over 30, 60, 90, 120, or 180 minute gradients running from 96% Buffer A (5% acetonitrile, 0.125% formic acid) and 4% buffer B (95% acetonitrile, 0.125% formic acid) to 30% buffer B. Sample eluate was electrosprayed (2,700 V) into a Thermo Scientific Orbitrap Eclipse or Orbitrap Ascend mass spectrometer for analysis. The scan procedure for MS1 scans (Orbitrap scan at 120,000 resolving power, 50 ms max injection time, and AGC target set to 100%) and MS2 scans (linear ion trap, “rapid” scan rate, 50 ms max injection time, AGC target set to 200%, CID collision energy of 35% with 10 ms activation time, and 0.5 m/z isolation width) was constant for all analyses with a gradient length above 30 minutes. For 30 minute gradient methods, the ion trap scan rate was set to “turbo” and the maximum injection time lowered to 11ms. Peaks from the MS1 scans were filtered by intensity (minimum intensity > 5e3), charge state (2≤z≤6), and detection of a monoisotopic mass (monoisotopic precursor selection, MIPS). Dynamic exclusion was used, with a duration of 90s, repeat count of 1, mass tolerance of 10ppm, and with the “exclude isotopes” option checked. High field asymmetric waveform ion mobility spectrometry (FAIMS) was set at “standard” resolution, 4.6 L/minute gas flow and 3 CVs: −40/−60/−80. Each CV value was set to top N mode with number of dependent scans set to 6. For MS3 scans, SPS ions were set to 10, MS1 Isolation window was 2 m/z and MS2 isolation window was 3 m/z. MS3 scans were performed at a resolving power of either 45,000 (Ascend) or 50,000 (Eclipse) with an HCD collision energy of 45%.

For sequential RTS/RTLS analysis, a single CV of −50 was used. MS2 nodes are set up sequentially with a real-time filter after each scan, the first filter is set to “trigger-only” and the filer set to pass along every MS2 (Cosine Score less than 100/minimum XCorr of 0) and the second filter set to block any further scans (Cosine Score/XCorr greater than 100). For FTMS2 analysis in this experiment, a resolving power of 15,000, a max fill time of 54ms and an AGC target of 200% were used. All raw data are available through accession PXD039855.

### Real-time spectral library searching

Predicted spectral libraries were generated using Prosit-TMT (https://www.proteomicsdb.org/prosit/) and output to ‘.msp’ files. MSP files were converted to an RTLS/mzVault compatible format (‘.db’ extension) based on an indexed SQLite database structure. These files can be consumed by the RTLS filter in the instrument’s method editor. Conversion was done using the in-house developed DBKey R Shiny application (https://github.com/SchweppeLab/DBKey). Unless otherwise noted, libraries were generated with no missed cleavages, fixed TMTpro at n-term and lysine and variable oxidation on methionines. For each library candidate within the specified precursor tolerance, a cosine similarity score was calculated against the acquired spectra, with the cosine similarity score defined as such (Equation 1):

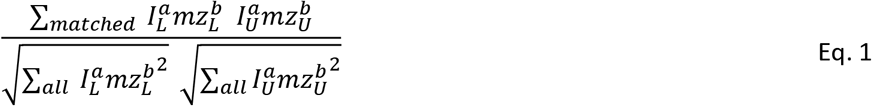

where *L* is library peaks, *U* are acquired spectral peaks, *I* is intensity of peak, *mz* is m/z of peak, *a* and *b* are the weight factors. Unless otherwise specified, the minimum cosine score threshold used was set to 20 approximate the triggering rate of RTS-based methods.

Real-time spectral searching and analysis was done using Orbitrap Eclipse instrument control software version 4.0, unmodified from the released version except where noted. Unless otherwise specified, the minimum cosine score threshold used was 20. Precursor tolerance was set to 10 ppm and isotope correction was set to 0/1. TMT-SPS mode was selected, such that only matched fragments with a TMTpro tag were selected for SPS-MS3 scans. For methods targeting yeast in HYPro, peptides derived from human proteins were rejected from triggering further SPS-MS3 scans, using the keyword promote/reject feature. Spectral libraries built with SpectraST 5.0 were processed through the Trans-Proteomic Pipeline^33^. Raw files were converted to mzXMLs with msconvert and subsequently searched with Comet using settings described in the data analysis section^34^. Results were then processed using PeptideProphet before SpectraST was used to build a consensus library with default settings. Decoy library entries were generated via precursor swap ^35–37^.

### Real-time database searching

Real-time search was done using the instrument control software, Tune version 4.0, unmodified from the release version except where noted. For the fractionated HDAC-treated cell line experiment, a human database was used with fixed C(Cam) and variable M(Ox), one missed cleavage, with TMT mode enabled, a minimum XCorr of 1.0, a minimum dCn of 0.05, precursor tolerance of 10 ppm and FDR/protein closeout disabled. Unless otherwise noted for HyPro runs, a concatenated human-yeast database (a single FASTA file starting with the human proteome then the yeast protoeme) was used with the same modifications, “TMT mode” enabled, a minimum XCorr of 1.4, a minimum dCn of 0.05, and precursor tolerance of 10 ppm.

### Post hoc data analysis

Raw files were searched using the Proteome Discoverer 3.1 software. Unless otherwise noted, Comet and Sequest searches were performed against databases downloaded from Uniprot, with 2 missed cleavages and a 20 ppm precursor mass tolerance. For ITMS2-MS3 methods, a 0.6 Dalton fragment tolerance was used, and for FTMS2 methods, a 10 ppm fragment tolerance was used. Charge state was restricted to 2-6 and peptide length was limited to 7-30 amino acids to comply with INFERYS. All searches were performed with variable methionine oxidation (+15.99491), static cysteine carboxyamidomethylation (+57.02146), and static TMTpro modifications on lysine and the peptide N-termini (+304.207126). MSPepSearch was run with an INFERYS predicted library with 1 missed cleavage, fragment charges 2 to 4, variable methionine oxidation (+15.99491), static cysteine carboxyamidomethylation (+57.02146), and static TMTpro modifications on lysine and the peptide N-termini (+304.207126). MSPepSearch precursor tolerance was 10 ppm and the fragment tolerance was 0.6 Dalton for ITMS2-MS3 methods and 10 ppm for FTMS2 methods. Peptide spectral matches and spectrum spectral matches were filtered to a peptide and protein false discovery rate (FDR) of less than 1%^38^. Quantification was done by selecting the centroid with the highest signal-to-noise ratio within a 0.003 Dalton tolerance of the reporter ion’s theoretical m/z. Unless otherwise noted, peptides identified using FTMS2 or SPS-MS3 methods were considered quantified if the sum of the reporter ions’ signal-to-noise (“s:n”)ratios was greater than 100 and precursor isolation specificity was greater than 0.5. Peptides identified using RTS or RTLS methods were considered quantified if the sum of the reporter ions’ signal-to-noise ratios was greater than 100, regardless of precursor isolation specificity. Chimeric spectra analysis was performed using CHIMERYS under the same settings as above, using the “inferys_2.1_fragmentation” prediction model. Quantitative accuracy was assessed with the HYPER interference-free index (IFI, Equation 2), a modified version of the original TKO interference free index where the empty channels in the HyPro standard serve as substitute for the empty KO channels ^39,40^.

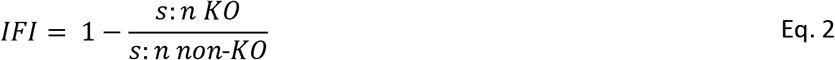

Statistical analyses and plotting was done using the R project for statistical computing^41^.

## Results and Discussion

Spectral library searching has been shown previously to increase the sensitivity of peptide detection in quantitative proteomics by leveraging either acquired spectra or predicted fragment ion intensities^42,43^. With a diverse array of scoring metrics, analysis pipelines, and applications, spectral library searching has proved to be a robust method for a wide array of quantitative proteomics methods, though it is predominantly used for label-free quantitation, and data-independent acquisition methods (DIA)^44^. To enable spectral library searching for real-time decision making we had to (1) determine a method and memory compatible means to store full-proteome spectral libraries (2) enable conversion of diverse spectral library formats into this common format, and (3) optimize scoring functions for performant real-time decision making (Figure 1A).

**Figure 1.**
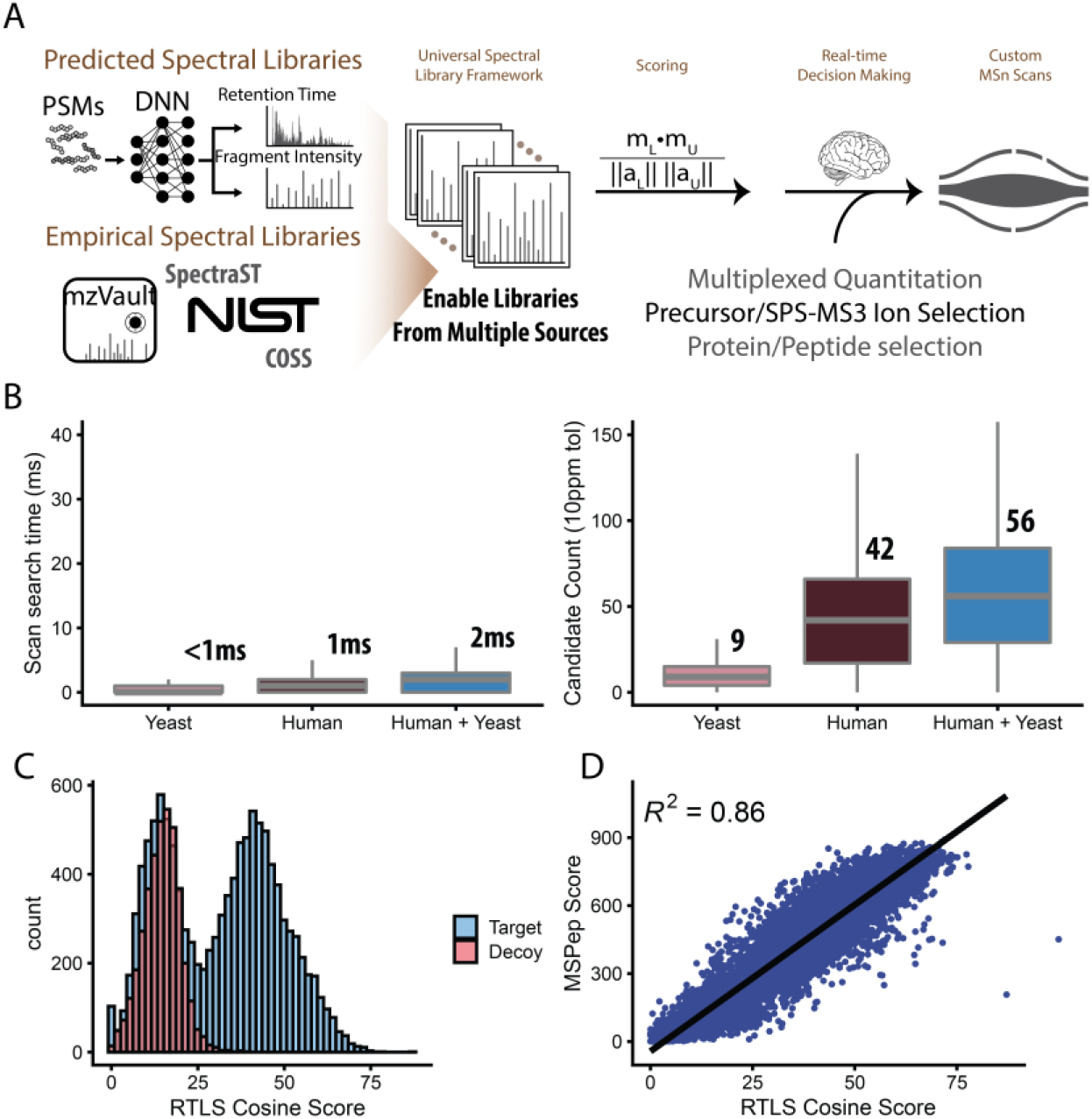
Overview of RTLS workflow. A) The DBkey-RTLS workflow enables the use of spectral libraries from repositories, deep neural network (DNN) predictions, and empirical data. Spectral library matching and scoring are then used to define subsequent scans (SPS-MS3). B) (Left) Boxplots of the search time of Prosit-TMT libraries for yeast, human, concatenated human-yeast. (Right) Library candidates scored per scan for the same libraries. C) Target-decoy separation for a yeast sample labelled with TMTPro. D) Scatterplot comparing RTLS cosine score to that of an established offline spectral library search, MSPepSearch.

We began by building a library processing tool, called DBKey, that can take in spectral libraries from both empirical and predicted spectral sources including repositories such as NIST, SpectraST, and Prosit. DBKey then converts these input file types to a single, RTLS-compatible data type for real-time processing. Our common data type is also compatible with mzVault and is based on an indexed SQLite data structure that can be held in memory during instrument acquisition and stored for repeated use (‘.db’ file extension). By integrating DBKey into a Docker image, we can process these files efficiently in a Docker environment for workstation or cloud extensible deployment.

Using DBkey, we built libraries from both predicted libraries and publicly available datasets to evaluate their compatibility with a multiplexed RTLS-MS3 workflow. Prosit-TMT^19,20^ was used to make whole-proteome predicted libraries from *S. cerevisiae* and human FASTA files (497,718 and 2,225,832 spectra respectively) (Figure 1B). Queries to the database generally require less than 1ms for single proteome spectral libraries even when the library consists of millions of individual spectra (Figure 1B).

While cosine score weights have been optimized previously for label-free proteomes^45^ they had never been tuned for TMTpro-labelled peptides. We performed a parameter sweep for the scoring based on Equation 1 and found that the weights *a* = 0.4 (applied to the intensity) and *b* = 0.9 (applied to the m/z) provided the highest sensitivity for peptide spectral matching of TMTpro labeled peptides (Figure S1). This was in-line with previous work that found down-weighting dominant peaks and up-weighting larger fragment ions improves sensitivity^30^. PSMs scored with our optimized cosine score discriminate target and decoys well and show strong agreement with Comet database search results (Figure 1C).

Using the optimized score weights, we sought to establish RTLS’ potential for efficiently matching spectra in real time with instrument acquisition. Ideally, spectral matching should occur fast enough to enable parallelization scan acquisition. Using optimized indexing from the .db files, we were able to match experimental spectra to library spectra from predicted and empirical libraries with median search times of 1 ms for single proteome databases (human or yeast) and 2ms for a concatenated human-yeast library (Figure 1B). Importantly, even as the number of candidate spectra considered for each search increased, the relative search time remained well below our target of 35ms. RTLS maintained accurate discrimination of target and decoy peptides using the weighted cosine score (Figure 1C) and these scores correlated well with post hoc spectral match scoring from MSPepSearch (Figure 1D, Figure S2C).

While we performed our initial post hoc analysis with MSPepSearch, we wanted to evaluate multiple search pipelines to determine the optimal post-RTLS methods for sensitive detection of labeled peptides. To this end, we searched a 12-fraction whole-proteome multiplexed sample set collected with RTLS methods with three different search workflows: Comet (canonical database searching), MSPepSearch (spectral library searching), and Sequest with INFERYS^23^ rescoring – a database search rescored on spectral similarity to predicted fragment ion intensities (Figure S2). Settings were kept as similar as possible across informatics workflows (see methods). Though we observed a high degree of overlap between search methods, Sequest+INFERYS returned the highest number of confidently identified PSMs (Figure S3). Due to the improved detection sensitivity for PSMs and the incorporating of aspects of database and library searching with Sequest+INFERYS, we proceeded to use this pipeline for post hoc analysis of RTLS acquired data.

Spectral library searching is a highly flexible approach, but can be influenced by library spectra sourcing, peptide fragmentation, spectral quality and spectral purity during instrument acquisition^46^. To test this, we measured peptide detection sensitivity and quantitative accuracy across a panel of acquisition methods comparing: (1) predicted or empirically derived spectral libraries, (2) with or without FAIMS, and (3) using both CID and HCD MS2 fragmentation (Figure S5-S7). First, we generated an empirical spectral library from fractionated yeast samples labeled with TMTpro (PXD014546)^47^ and assembled a library using SpectraST (66,415 spectra). We compared this empirical yeast library to a predicted library built using Prosit-TMT. Using these two libraries we found that RTLS could efficiently match spectra and trigger SPS-MS3 scans for both empirical and predicted libraries. We observed the cosine scores distribution skewed higher in the empirical spectra results, most likely due to incorporation of non b/y fragments, yet the number of quantified peptides and the concordance with offline searching was lower (Figure S5). Thus, when using empirical libraries, it is important to determine score distributions to establish an optimal score threshold for a given filter-library pairing. While scoring for an empirical library would likely be improved using data generated on the same instrument and optimized library construction, the promising results, robustness, and flexibility of predicted libraries, we primarily used Prosit-derived libraries for this work.

Second, when comparing RTLS method s with or without using FAIMS, we observed a broader score distribution, higher median score (20.7 to 26), and better target/decoy discrimination (Figure S6). We believe this is due to FAIMS ability to reduce precursor co-isolation^40^ thereby generating spectra with fewer interfering peptide fragments and higher spectral match scores. For this reason, we proceeded with FAIMS for further experiments. Third, peptide fragmentation methods used during acquisition can greatly influence library-spectral matching^11^. We examined both CID and HCD RTLS workflows for use in multiplexed method. For these analyses, we tested if predicted library spectra based on a specific fragmentation type and energy would affect the sensitivity of RTLS peptide detection (Figure S5). Due to the similar score distributions and small (3.8%) difference in peptides detected we chose to use CID peptide fragmentation as it is the most similar to canonical SPS-MS3 workflows (Figure S7).

Having established an optimized set of methods parameters we began by comparing RTLS to traditional SPS methods for whole proteome single-shot methods. Running 180 minute gradients with HyPro we confirmed that RTLS resulted in more selective triggering and an increase in quantified peptides and proteins of 26.8% and 15.8%, respectively (Figure 2A). We then moved to the challenging task of sub-proteome analysis, quantifying only a subset of our sample, in this case the lower-abundant yeast proteins in our standard samples (10% of the peptides by mass). When comparing SPS-MS3 to RTLS methods of equal length (180 minute gradients) RTLS increased the number of unique quantified peptides and proteins by 42% and 64.4%, respectively. When RTLS methods targeting yeast proteins were shortened to only half of the gradient time (90 minute) of the SPS-MS3 methods, RTLS still quantified 3.1% more unique peptides and 15.9% more proteins (Figure 2D). We believe that the stronger improvement for proteins quantified was due to sampling of single peptides that were only captured because of RTLS methods are able to sample more precursors to generate, search, and score more MS2 spectra. When we compared the fraction of MS3 spectra triggered during instrument acquisition were used to obtain our final set of quantified peptides and proteins, we found that RTLS increased MS3 usage from 4.5% to 74.8% (Figure 2E).

**Figure 2.**
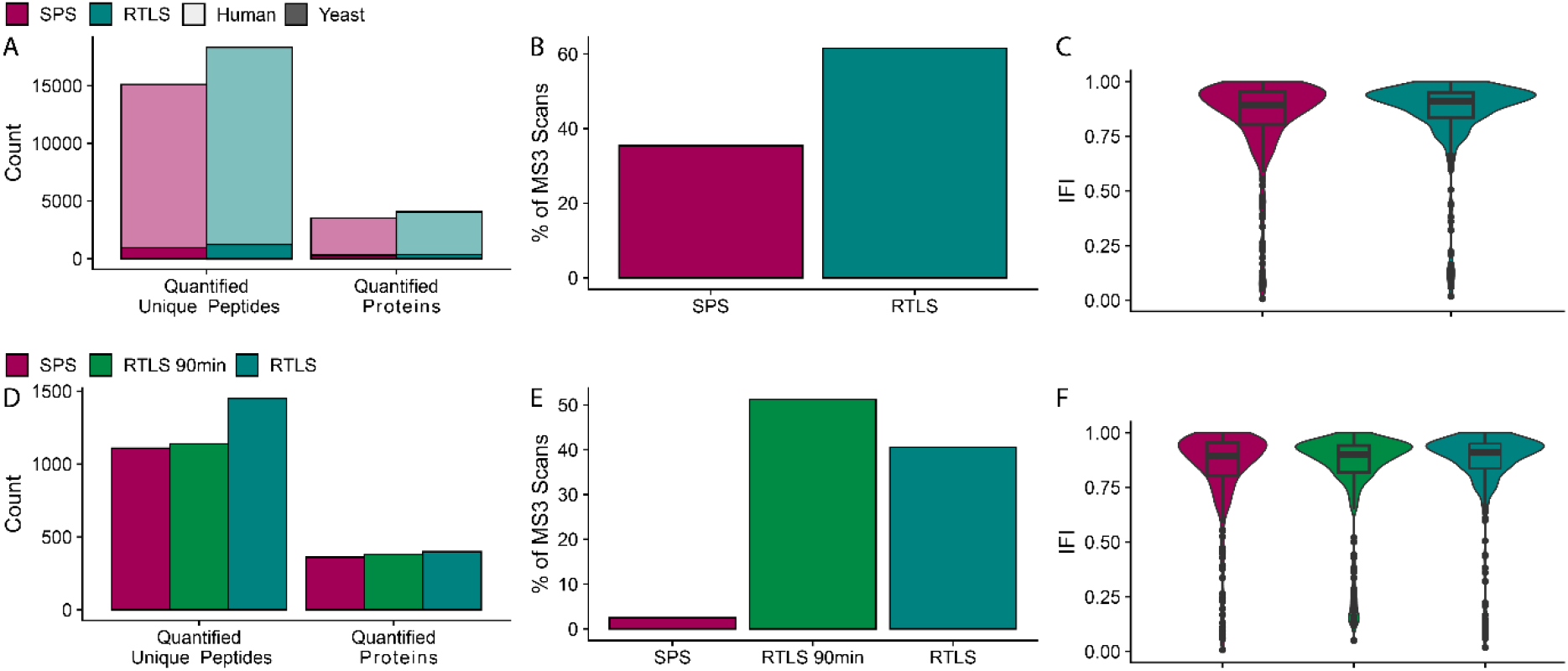
Quantifying peptides and proteins from human and yeast in HyPro. A) Quantified peptides and proteins from 1ug HYPro standard in single-shot 180 minute runs using either SPS or RTLS. B) Percent of total acquired MS3 scans that led to quantified peptides included in the final results C) Violin plot showing the HYPER interference-free index of Yeast proteins quantified.

We next sought to investigate whether one or the other of these methods was better suited to sample multiplexed proteomics analysis. To do this, we ran a series of single-shot methods with settings matched as closely as possible between RTS and RTLS so as to generate similar rates of triggering quantitative MS3 scans (Figure 3A). We observed a slight but consistent increase in the number of quantified peptides when using the RTLS methods (Figure 3A). To determine why RTLS consistently generated more quantified peptides we ran a sequential real-time method to directly compare RTS and RTLS on the same set of precursors in the same analytical run. In this method, the MS2 level was branched so that each precursor produced two MS2 spectra. Each of these subsequent MS2 scans were then analyzed in real time by either RTS or RTLS to generate a matched set of filtering and quantification events. We then processed this data through the common Sequest+INFERYS pipeline which combines database searching and library rescoring. With the matched set of RTS and RTLS triggering events, we compared XCorr (Comet-based RTS) and weighted cosine score (from RTLS) for their ability to classify target and decoy peptides (Figure 3B). In replicate analyses, we found that RTLS was more sensitive at low FDR thresholds (1-5%) at detecting confirmed peptide spectral matches from the post hoc analysis.

**Figure 3.**
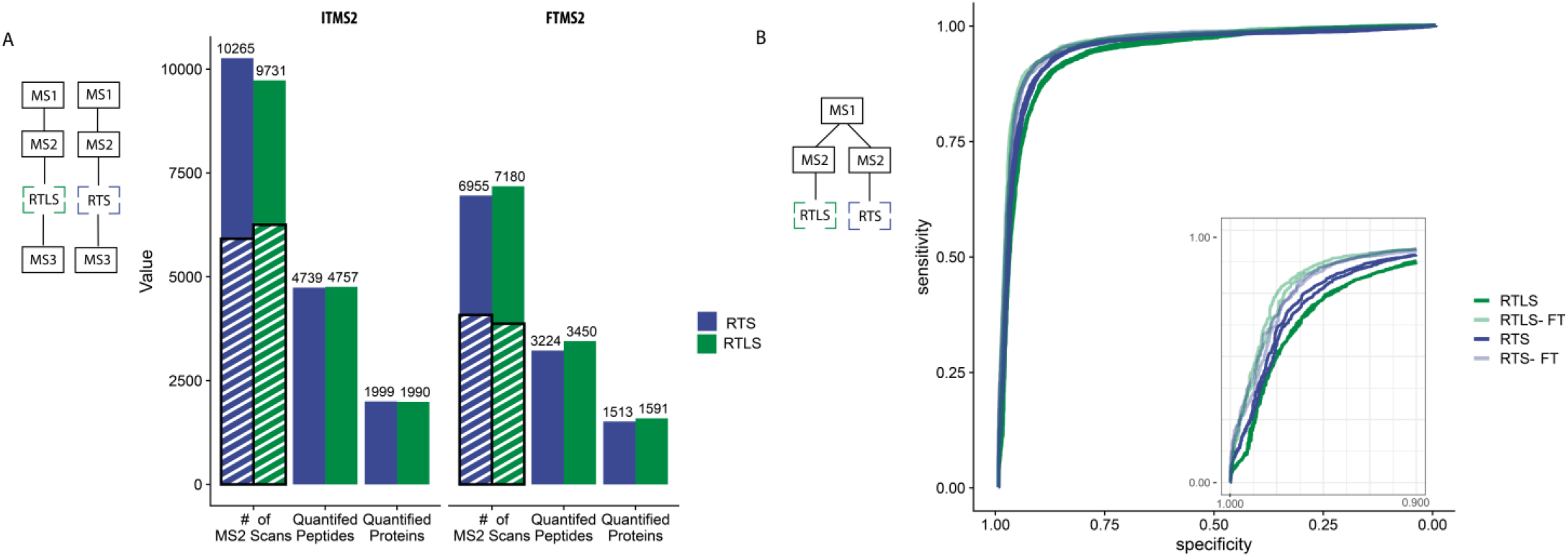
Comparing IDA methods. A) 30 minute single shot runs of HyPro comparing RTS and RTLS done with MS/MS from both the ion trap (IT) and Orbitrap (FT). MS2s, MS3s, quantified peptides and quantified proteins plotted. MS3s are represented by overlaid stripes. B) Receiver-operator characteristic plots of both ion-trap and Orbitrap MS2 spectral matching with either RTS or RTLS based on XCorr (RTS) or Cosine Score (RTLS).

While single-shot methods are valuable, one of the most common uses for multiplexed proteomics is for the analysis of fractionated proteomes, quantifying differential abundance across conditions. To test RTLS methods across fractionated samples, we treated three human cell lines (A549, H292, PSC1) with the histone deactylase inhibitor belinostat, using four different acquisition methods: SPS-MS3, HRMS2, RTS, and RTLS. Perturbation with belinostat treatment alters chromatin state and leads to pleotropic remodeling of the proteome after sustained treatments. In keeping with previous analyses, we compared belinostat treatment responses with SPS samples run as twelve 180 minute runs while the HRMS2, RTS, and RTLS methods were collected using half of the gradient time (90 minutes per fraction). We found that both RTS and RTLS methods quantified 4-15% more significant differentially abundant proteins (q-value < 0.05, |log_2_ fold-change| > 1) across all three cell lines when compared to the SPS-MS3 method (Figure 4A). Comparing the log_2_ fold-changes of shared significantly quantified proteins, IDA methods have higher absolute fold-changes due to SPS ion selection of b- and y-ions leading to improved quantitative accuracy (Figure 4B, Figure S8).

**Figure 4.**
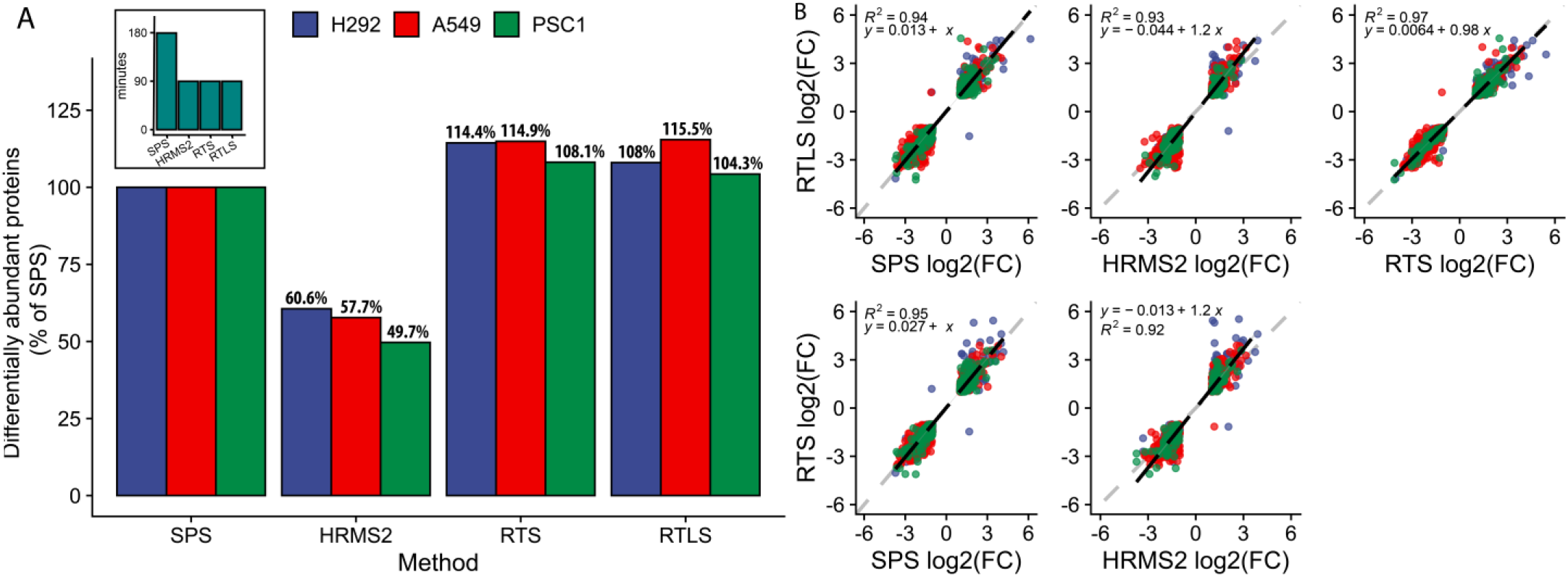
Use of RTLS in fractionated whole-proteome experiment. A) Differentially abundant proteins (| Log2FC| > 1 and q-value < 0.05) for H292, A549, and PSC1 cell lines as a percentage of the SPS results. (Inset) analysis time per fraction B) Scatter plots comparing the log2 fold changes of common significantly differential proteins found between different methods with the correlation and slope of linear regression.

We extended our study of whole proteome perturbations to compare determine the benefits of RTLS when using new instrumentation for sample multiplexed quantitative proteomics workflows. In particular, we wanted to determine if improved quantitation of peptides when running RTLS methods. Therefore, we compared the reporter ion sensitivity for matched RTLS methods and matched LC systems to quantify the proteomes of A549 cells treated with three HDAC inhibitors (abexinostat, CUDC-101, vorinostat). From fractionated analysis of short gradient runs (60 minute) on both an Orbitrap Eclipse and Orbitrap Ascend using RTLS, we found that the Orbitrap Ascend increased the detected reporter ion signal-to-noise by 127% for peptides and 145% for proteins when running RTLS methods (Figure S9).

Including the work above, most multiplexed proteomics methods are designed to minimize or mitigate precursor co-isolation, as failure to do so leads to chimeric spectra, which impair accurate quantitation. However, co-isolation remains a general challenge in complex sample analyses. Interestingly, recent work with library searching of label-free samples illustrated that purposefully generating chimeric spectra can increase identifications^48^. We hypothesized that if RTLS can correctly identify multiple precursors in a chimeric MS2 spectra, we could potentially trigger multiple separate MS3 scans from the same MS2 that would lead to an increase in sensitivity and quantified peptides across the run. To this end, we tested methods where the MS2 isolation width was increased from 0.4 Th to 2.0 Th and enabled the multiple precursor search in RTLS. This option allows the search engine to consider multiple precursors within the isolation window when performing the search on a single MS2 spectrum.

In developing multi-precursor RTLS methods, we found that canonical post hoc searching could not efficiently identify peptides from chimeric spectra. To address this, we performed post hoc searching using CHIMERYS, a newly developed search engine focused on deconvolution of chimeric spectra^49^. CHIMERYS successfully validated hundreds of chimeric spectra from our multiple precursor methods. Importantly, we found that RTLS could also correctly identify multiple precursors from a single MS2 scan. As a proof of principle, CHIMERYS identified 77.9% of all PSMs to be derived from chimeric spectrum when using a wide 2.4 Da isolation width (Figure S10). In 43.6% of the chimeric spectra, RTLS returned at least one of the CHIMERYS validated peptides as the top precursor match. There was concordance between RTLS and CHIMERYS on multiple PSMs within a spectrum across 13.5% of the validated chimeras. Due to the differing known quantitative profiles of the human and yeast peptides in our HyPro standard we were able to validate detection of chimeric peptide spectra from a single MS2 scan.

We found that RTLS could properly assign both a human and yeast peptide to a chimeric MS2 spectra and then trigger MS3-based quantification consistent with the known concentrations across TMTpro channels (Figure 5C). We calculated interference free indices (IFI) for both the human peptide (IFI = 0.12) and yeast peptide (IFI = 0.89) and observed distinct quantification profiles across TMTpro channels that were consistent with the coisolation and fragmentation of two peptides (Figure 5C). Future improvements in methods for chimeric spectra generation and detection sensitivity are still needed to implement this method. Despite these considerations, chimeric spectra triggering with RTLS serves as the first step towards addressing chimeric spectra isolation in sample multiplexed proteomics and potentially leveraging wider isolation width methods for isobaric multiplexed samples.

**Figure 5.**
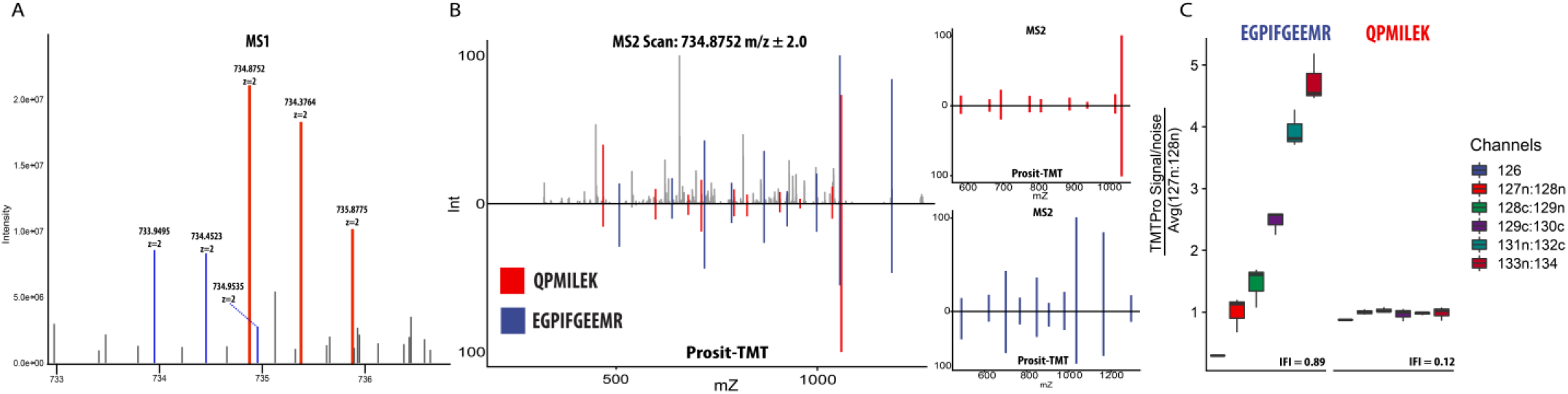
Chimeric spectra and RTLS. A) Acquired MS1 scan with overlapping precursor isotopic envelopes. B) ddMS2 triggered from the MS1 in A with matched RTLS fragments colored for both precursors (right) Mirror plots of the fragments matched by RTLS and the library entry from a concatenated Prosit library of human and yeast peptides. C) Hypro interference free indexes (IFI) of the two MS3 spectra that were triggered from the single MS2 spectrum in panel B. Channels are grouped by their expected ratios (see Materials and Methods).

## Conclusions

We have reported the development and use of RTLS, a modular, integrated intelligent data acquisition strategy based on spectral library searching in real time for sample multiplexed proteomics. The use of RTLS methods resulted in improved instrument efficiency and increased the number of proteins and peptides quantified when compared to traditional methods. Establishing RTLS for multiplexed proteomics lays the groundwork for future work utilizing library searches in areas where library searching can potentially make a large impact. This includes studies of post-translational modifications, where including modifications in the match scoring can lead to large search spaces. As we have shown, RTLS will also be useful in chimeric spectra deconvolution, and in combination with other IDA methods, such as RTS, for highly selective and adaptive instrument methods. In addition to the core methods, we present optimized RTLS scoring weights for sample multiplexed analyses and demonstrate the utility of integrating RTLS and FAIMS for improved discrimination of low confidence peptides. These optimizations can further be improved upon using new instrumentation to increase the sensitivity of detection of quantified peptides and proteins. Finally, we demonstrate that RTLS is capable of triggering multiple, quantitatively distinct MS3 spectra from the same MS2 spectrum. Together these findings highlight how new IDA methods can be used to improve sample multiplexed quantitative proteomics methods.

## Acknowledgements

We would like to thank the Schweppe Lab and Jimmy Eng of the UWPR for helpful advice.

