## Supplemental Information for "Real-time spectral library matching for sample multiplexed quantitative proteomics"

**Contents**

*Figures*

Figure S1. Optimizing cosine score for TMT labelled peptides.

Figure S2. Scatter plots for RTLS and other scores.

Figure S3. Workflow for searching data.

Figure S4. Venn diagrams of search results.

Figure S5. RTLS libraries from SpecrtaST and Prosit.

Figure S6. Using FAIMS on RTLS.

Figure S7. Fragmentation method comparison.

Figure S8. Volcano plots from Belinostat experiment.

Figure S9. IDA methods on shorter gradients.

Figure S10. Percentage of chimeric MS2 spectra.

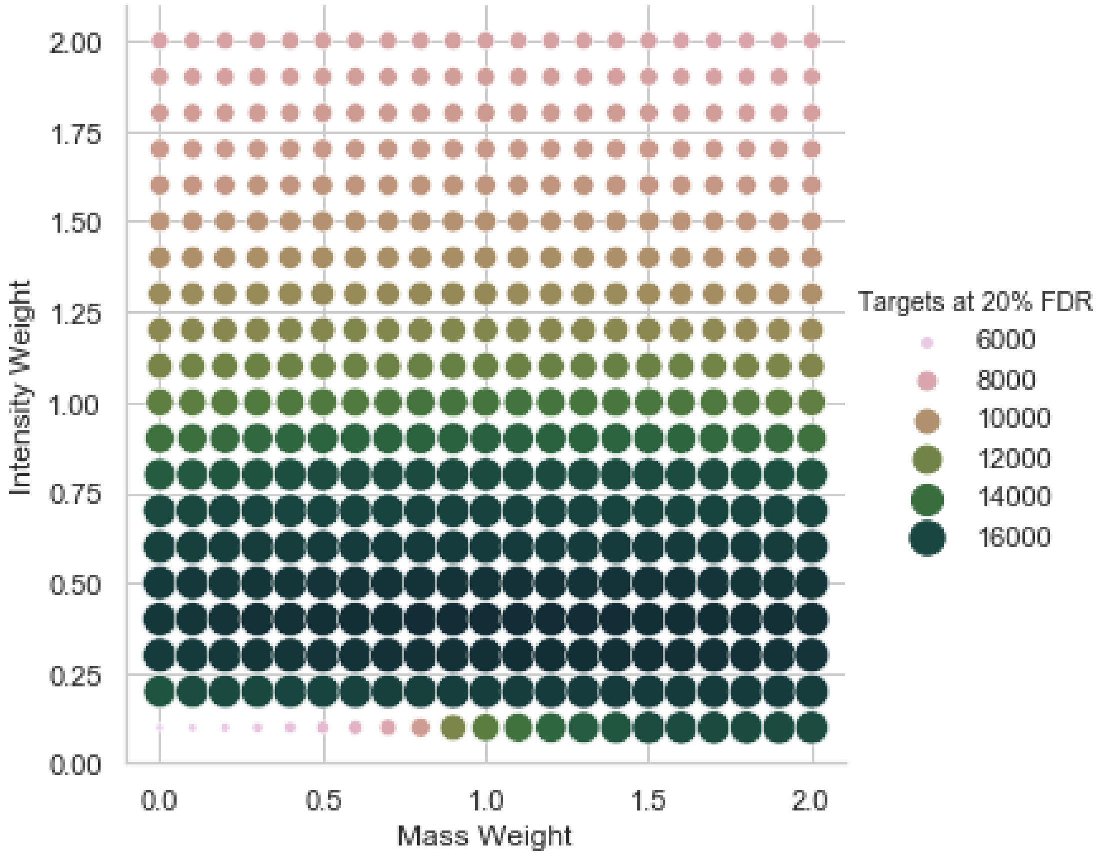


**Figure S1** Parameter sweep at different cosine score weights run on human and yeast TMTpro labelled peptides, measured at 20% FDR.

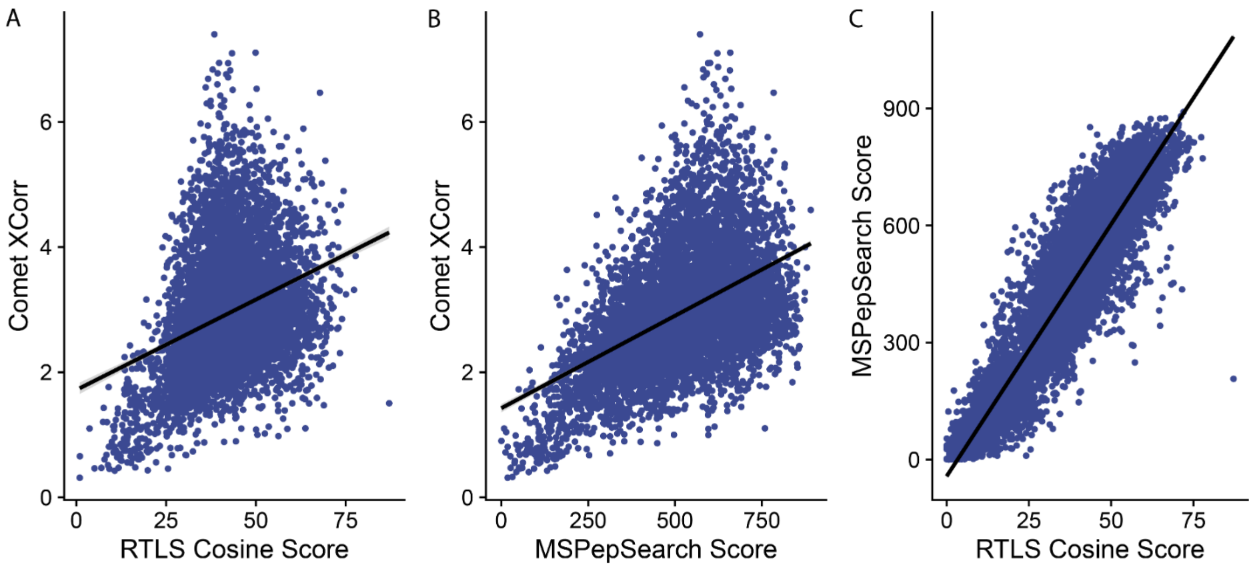


**Figure S2** Score comparisons for different proteomics search methods. A) Scatter plot of RTLS cosine score and Comet XCorr. B) Scatter plot of Comet XCorr and MSPepSearch Score. C) Scatter plot of MSPepSearch and RTLS Cosine Score.

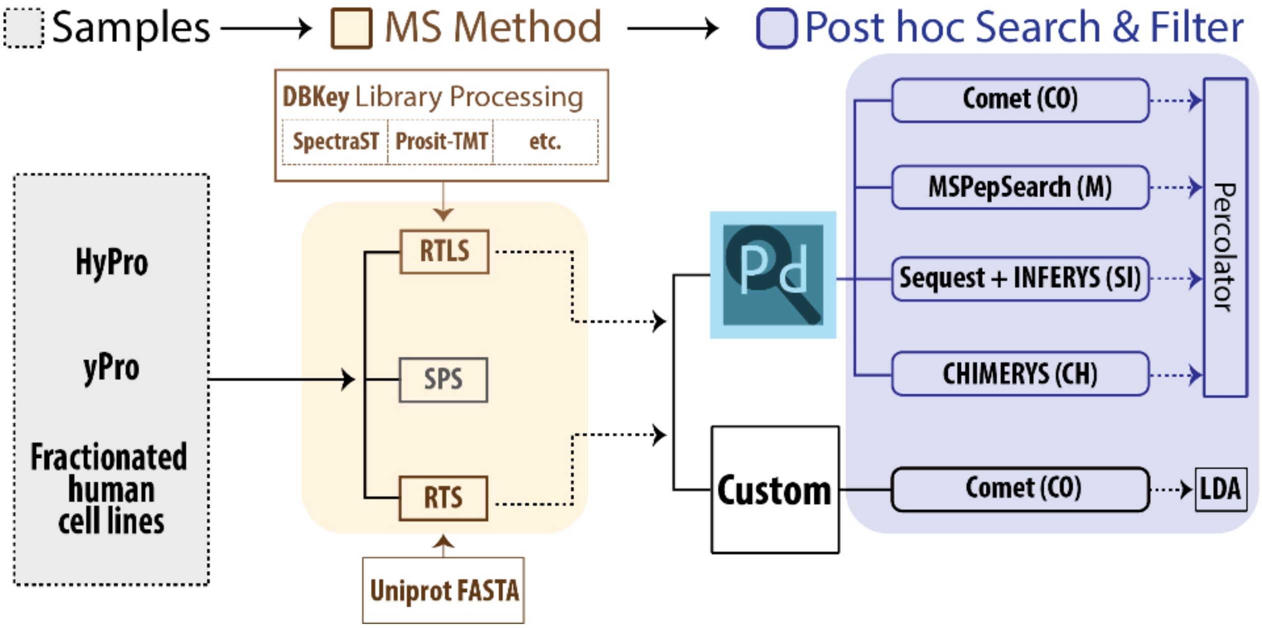


**Figure S3** Diagram showing the samples (HyPro, yPro, and fractionated human cell lines), methods (SPS, RTS, RTLS) and data analysis (Comet, MsPepSearch, Sequest+INFERYS, CHIMERYS) performed in this study.

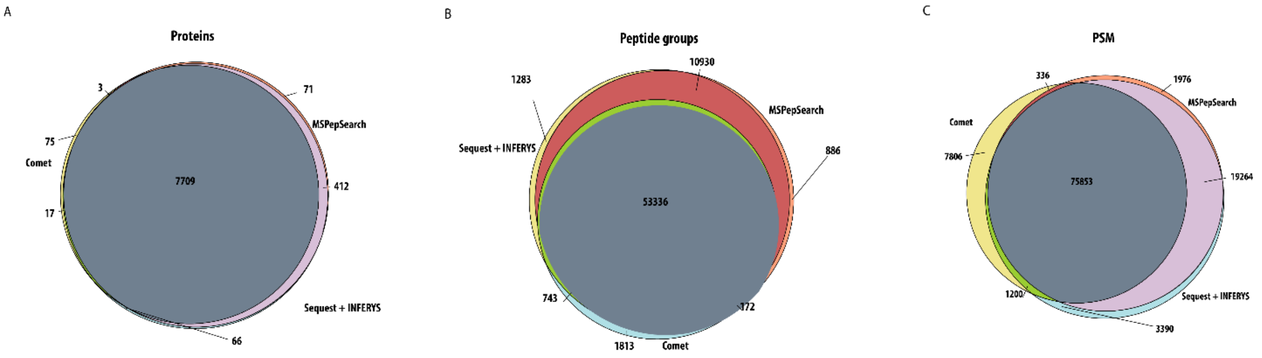


**Figure S4** Venn diagrams showing the overlap between Sequest + INFERYS, MSPepSearch and Comet in Proteome Discoverer at protein (A), peptide group (B), and PSM (C) level for fractionated whole proteomics data.

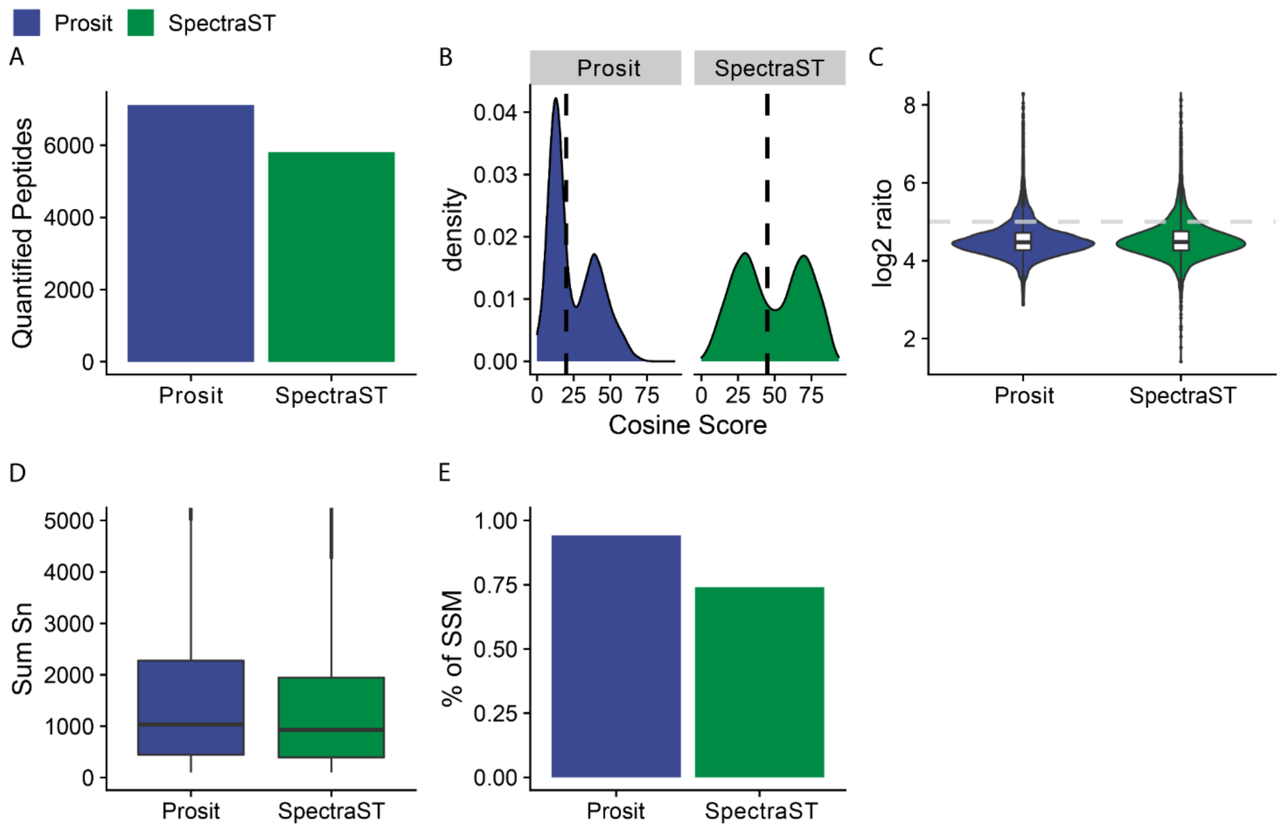


**Figure S5** Comparison between empirical and predicted spectral libraries. SpectraST libraries were built from fractionated DDA data with default settings. A) RTLS cosine score distribution. B) Violin plot showing quantitative accuracy by looking at the log2 ratio of 133n, 134c, 134n over 127n, 127c, 128n. Gray dotted line represents the median ratio when injected as single proteome. C) Percent of matches that agree with Comet database search. D) Comparison of PSM level reporter ion signal.

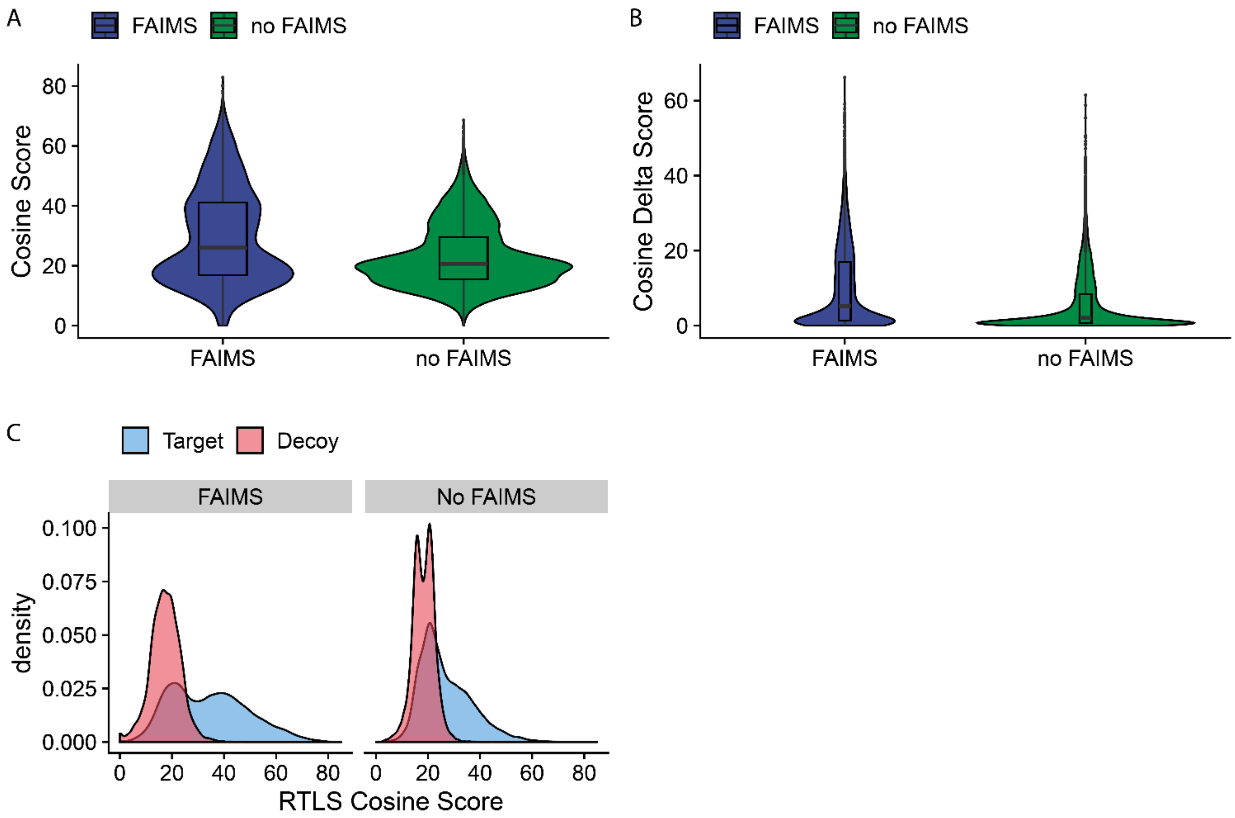


**Figure S6** RTLS with and without FAIMS. A) Violin plots showing cosine score distribution. B) Violin plots showing delta cosine score distribution. C) Target/decoy discrimination for FAIMS and no FAIMS.

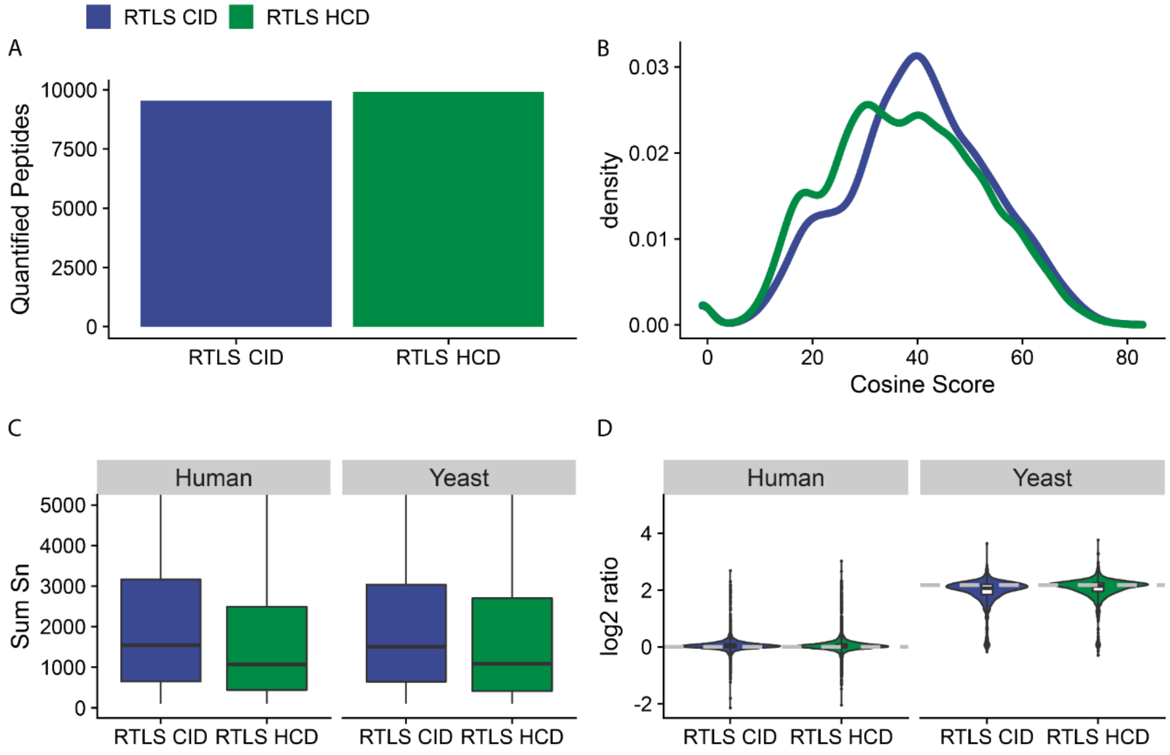


**Figure S7** RTLS run with HCD and CID collision energies. Prosit was used to generate CE specific library for HCD spectra. A) Comparison of quantified peptides in 120 minute run. B) Distribution of cosine scores between predicted library and acquired spectra. C) Boxplot comparing PSM reporter ion signal of the two proteomes found in HyPro. D) Violin plot showing quantitative accuracy by looking at the log2 ratio of 133n, 134c, 134n over 127n, 127c, 128n. Gray dotted line represents the median ratio when injected as single proteome.

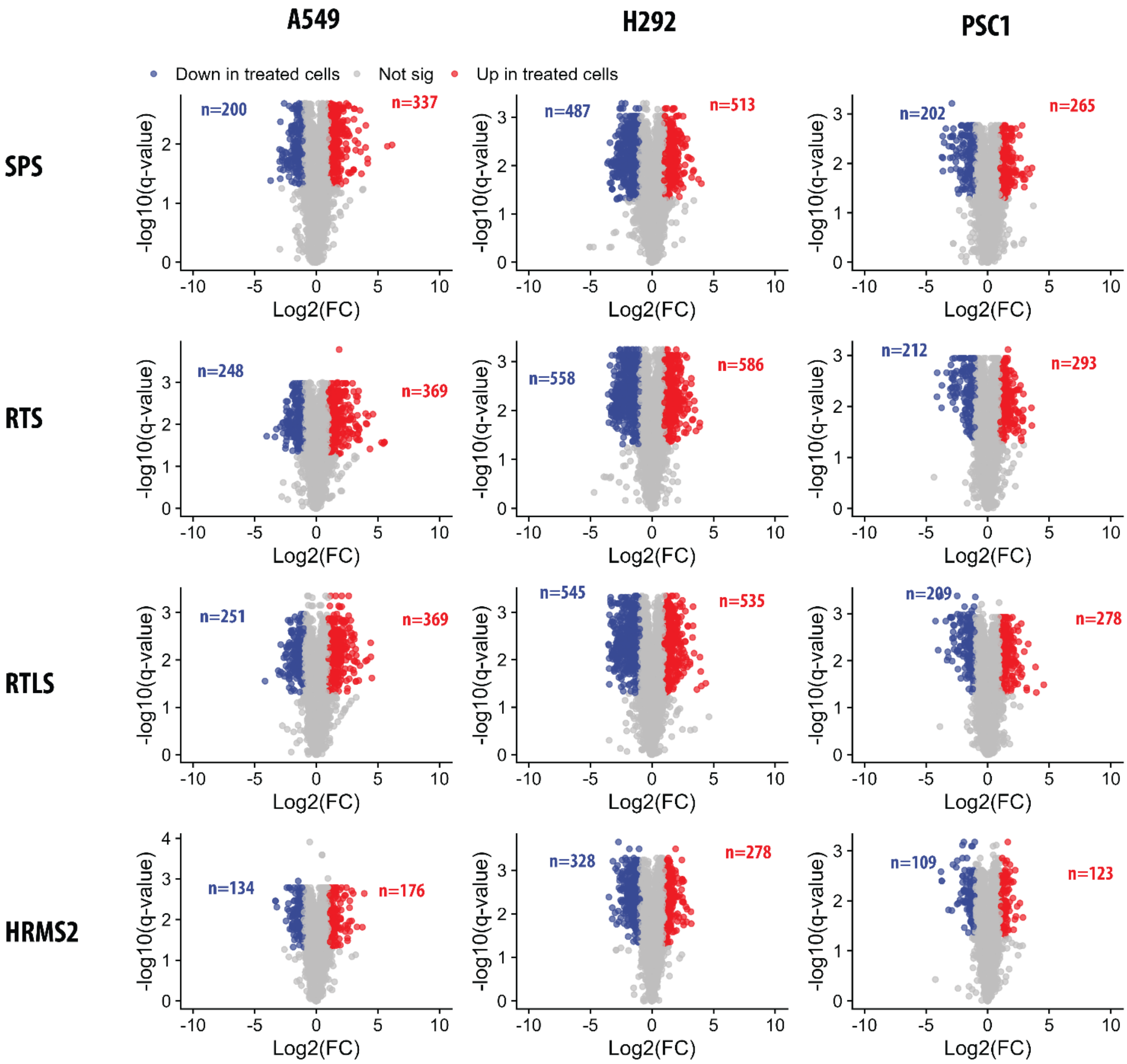


**Figure S8** Volcano plots from Belinostat experiment showing all three cell lines with the different acquisition methods. Proteins significantly more abundant than control are in red and less abundant in blue.

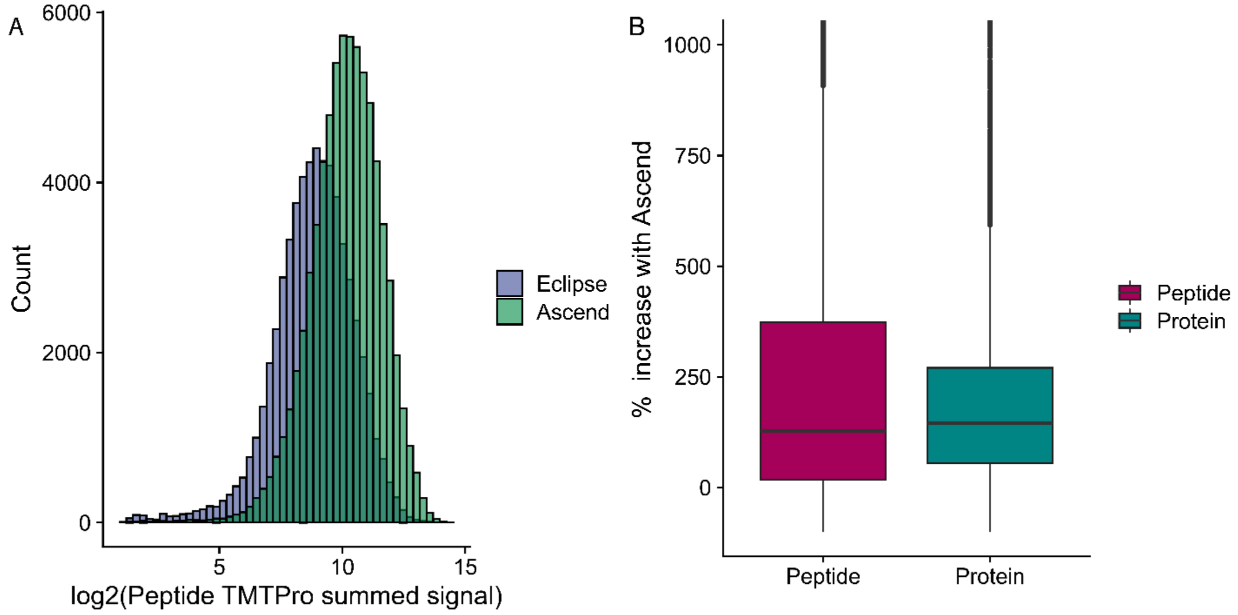


**Figure S9** A) Distributions of peptide level TMTpro signal for the two instruments on fractioned human cell lines. B) Boxplot showing percent signal increase for shared peptides and proteins.

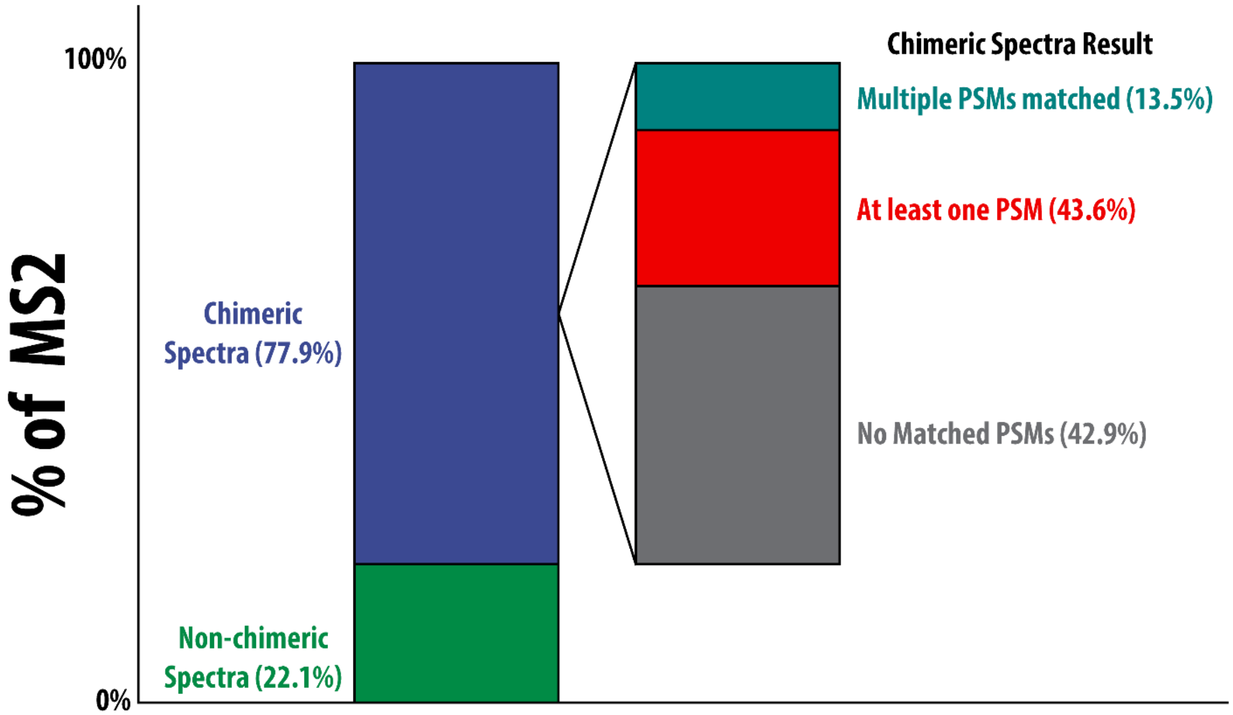


**Figure S10** Percentage of chimeric MS2 spectra. Spectra were collected from a 120 minute HyPro analysis using 2.4 Da isolation widths. Of the total MS2 spectra, 77.9% were determined by CHIMERYS to be chimeric spectra. Of the chimeric spectra, 13.5% were observed to match to multiple PSMs.
